# Inverted topographies in sequential fitness landscapes enable evolutionary control

**DOI:** 10.1101/2024.12.29.630702

**Authors:** Peng Chen, Nikhil Krishnan, Anna Stacy, Davis T. Weaver, Rowan Barker-Clarke, Michael Hinczewski, Jeff Maltas, Jacob G. Scott

## Abstract

Adaptive populations rarely evolve in a static environment. Therefore, understanding and ultimately controlling the evolution of a population requires consideration of fluctuating selective pressures. The fitness landscape metaphor has long been used as a tool for representing the selective pressures a given environment imposes on a population. Much work has already been done to understand the dynamics of evolution on a single fitness landscape. More recently, evolution on fluctuating or sequentially applied landscapes has come to the fore of evolutionary biology. As more empirical landscapes are described, metrics for describing salient features of paired landscapes will have uses for understanding likely evolutionary dynamics. Currently, Pearson correlation coefficient and collateral sensitivity likelihoods are used to quantify topographical relatedness or dissimilarity of a pair of landscapes. Here, we introduce the edge flip fraction, a new metric for comparing landscapes, which quantifies changes in the directionality of evolution between pairs of fitness landscapes. We demonstrate that the edge flip fraction captures topographical differences in landscapes that traditional metrics may overlook which have important consequences for the trajectories of populations evolving on them. By applying this metric to both empirical and synthetic fitness landscapes, we show that it partially predicts the collateral sensitivity likelihoods and can inform the optimality of drug sequences. We show that optimal drug sequences that keep populations within lower fitness regions require shifts in evolutionary directions, which are quantified by the edge flip fraction. Edge flip fraction complements existing measures and may help researchers understand how populations evolve under changing environmental conditions, and could yield clues in the pursuit of evolutionary control.

## Introduction

The ability to engineer and control evolution is a major outstanding question in biology with wide-ranging applications for human health [1], agriculture [2, 3], industry [4], and ecosystems [5, 6]. However, successful prediction and control remains difficult [7, 8]. Fitness (adaptive) landscapes, mathematical constructs that map an individual’s genotype to a phenotypic measure of fitness, have been proven to be a valuable model for diverse evolving systems, including bacteria [9], yeast [10], viruses [1], parasites [11] and cancer [12]. Over the last few decades, rapid experimental advances have made it increasingly possible to enumerate enormous genotype-to-phenotype maps, thus driving continued interest in theoretical modeling of fitness landscapes [13–19]. These experimental landscapes have revealed that mutations often have a complex relationship with the genetic background in which they arise (G×G interactions), their evolutionary environment (G×E interactions), or both (G×G×E interactions) [20, 21].

In particular, the topography of these fitness landscapes has sparked recent interest due to its role in influencing evolutionary trajectories [22, 23]. Critically, the topography defines how beneficial or deleterious emerging mutants are and therefore how likely they are to fix in the population. However, the topography of a fitness landscape is not static. Instead, the topography can fluctuate due to nonlinear interactions between genes (known as epistasis) [24–27], or environmental variation such as a varying antibiotic concentration [28], sequential drug application [29], or simply variable weather [30] and climate [6]. For example, collateral sensitivity — a phenomenon whereby a mutation that confers resistance to one environment also confers increased sensitivity to another unseen environment — would lead to fitness landscapes with negatively correlated topography [31–36]. In this context, a mutation that was once beneficial and fixed in one environment, is now deleterious and subsequently shed (or compensated for) in a following environment [37–39]. As a result, evolving populations are constantly challenged by fluctuating fitness landscape topographies that lead to dynamic fixation or extinction rates [40–48].

In this work, we seek to quantify the importance of this phenomenon — fitness landscape edge flips – where environmental variation leads to a dynamic fitness landscape topography that alters which mutations are beneficial or deleterious (and as a result “flips” the preferred direction of evolution between neighboring genotypes on the fitness landscape). We hypothesized that edge flips are critical to evolutionary control as they provide a genotype from which an experimentalist or physician may exert a particular selection pressure, favoring the selection of a mutant of their interest. We define a new metric for paired fitness landscapes the “edge flip fraction” which quantifies the extent to which two fitness landscapes preferred evolutionary trajectories differ. We show that the edge flip fraction is highly correlated with past metrics for paired fitness landscapes such as their Pearson correlation coefficient and collateral sensitivity likelihood. However, because edge flip fraction is sensitive to evolutionary transition probabilities themselves, rather than global structure like correlation coefficients, edge flip fraction proves to be a more valuable metric for identifying potential evolutionary control. To demonstrate this we leverage a previously published set of fitness landscapes to numerous *β* -lactam antibiotics [49] and reveal that the optimal sequence of antibiotics to minimize resistance enriches specifically for sequences of fitness landscapes with numerous edge flips. Finally, we show that because evolutionary control is leveraged at the genotypic level, it is the genotypic edge flip fraction (local edge flip fraction) that determines controllability.

## Results

#### Box 1: Edge Flip Fraction

**Figure 1.**
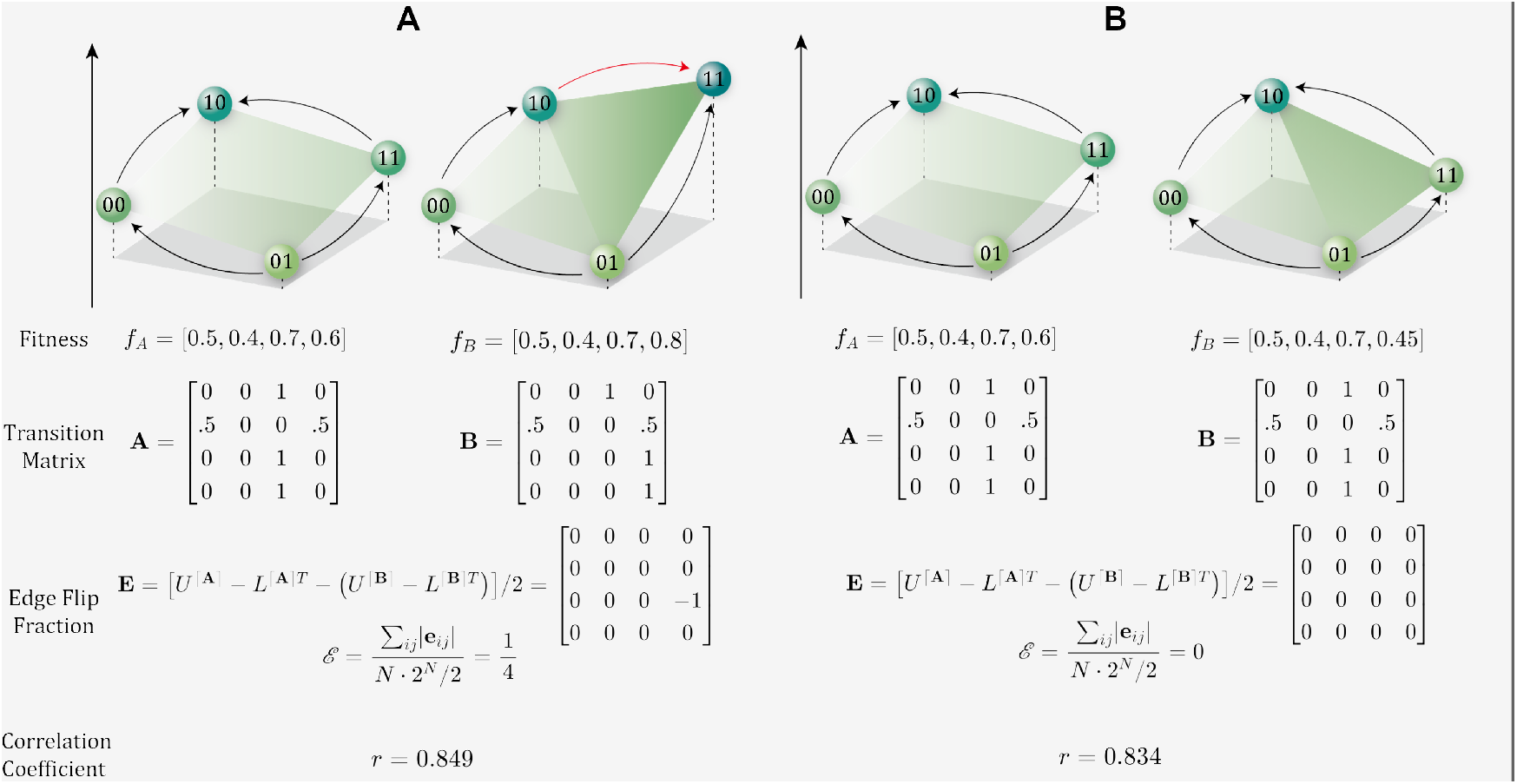
Two examples illustrating the edge flip fraction (ℰ ; see Methods for equation details). **A**. A pair of *N* = 2 landscapes in which increasing the fitness of the double mutant ‘11’ to 0.8, results in a Pearson correlation coefficient of *r* = 0.849 and leads to an edge flip, with an edge flip fraction of 0.25. As fitness of ‘11’ exceeds its Hamming distance-1 neighbor ‘10’, the direction of the edge between them changes. **B**. A pair of *N* = 2 landscapes with the same fitness values as in panel A, in which decreasing the fitness of the double mutant ‘11’ to 0.45 yields a similar Pearson correlation coefficient of *r* = 0.834, but does not result in an edge flip, with an edge flip fraction of 0. As long as the fitness of ‘11’ doesn’t go below that of its Hamming distance-1 neighbor ‘01’, the direction of the edge remains unchanged, so the changes in fitness are only reflected in the Pearson correlation coefficient.

### Edge flip fraction inherently contains information important for evolutionary dynamics

We consider the evolution of an asexual haploid population that can be characterized by a fitness landscape with *N* mutational sites with two alternative alleles (0 and 1). In such a model, each genotype can be represented by a bit-string of length *N*, which yields 2^*N*^ possible genotypes. We consider the evolution of this population in several environments (typically representing distinct antibiotics), where each genotype has a fitness value that corresponds to each unique antibiotic. In addition, we focus on small populations under strong selection (the so-called strong selection, weak mutation regime SSWM), whereby evolution can be characterized by random walkers on a fitness landscape. To illustrate the potential importance of edge flip fraction (ℰ ; see Methods), we begin by considering two pairs of *N* = 2 fitness landscapes with genotypes ‘00’, ‘01’, ‘10’, and ‘11’.

In the first landscape pair (**Figure 1A**), despite a relatively high correlation coefficient (*r* = 0.849), we observe an edge flip. Specifically, the difference in the fitness of genotype ‘11’ between the two environments leads to a change in the preferred direction of evolution along the connecting edge between genotypes ‘10’ and ‘11’. In contrast, the second landscape pair (**Figure 1B**) exhibits a comparable correlation coefficient (*r* = 0.834), but contains no edge flips. As a consequence, under the SSWM regime, the evolutionary trajectories are unchanged. This simple example highlights how edge flip fraction can capture important features important to the evolutionary dynamics on fitness landscapes that are overlooked by metrics like Pearson correlation coefficient.

To investigate this relationship in empirical fitness landscapes, we calculated the pairwise edge flip fraction and Pearson correlation coefficient across an ensemble of 15 four-locus fitness landscapes measured by Mira et al. [49]. Mira et al. characterized these landscapes by measuring the growth rates of 16 TEM *β* -lactamase genotypes—ranging from the wild-type genotype “TEM-1” through all possible combinations of four amino acid substitutions—in Escherichia coli treated with 15 different *β* -lactam antibiotics. The 15 drugs were categorized and color-coded by their classes, as detailed in **Table 1**. The edge flip fractions observed in the data ranges from 0.19 (CTX-CPD) to 0.72 (CTX-AM) (**Figure 2A**). Drug pairs from the same class, such as CTX and CPD (both classified as second generation Cephalosporin antibiotics), generally exhibit fewer changes in evolutionary direction, as evidenced by lower edge flip fractions. In contrast, cross-class drug pairs, such as CTX (Cephalosporin) and AM (Penicillin), tended to have higher edge flip fractions, suggesting that the broad biochemical mechanisms defining drug categories influence the topographical structure of fitness landscapes (**Figure S1**). Moreover, a comparison across all drug pairs revealed a moderate anti-correlation between edge flip fractions and Pearson correlation coefficient (**Figure 2B**). Intuitively, pairs of landscapes with few edge flips are also likely to be more correlated. These results suggest that edge flip fraction captures features of landscape topography similarly to the Pearson correlation coefficient.

**Table 1.** Name, class, and abbreviation for each antibiotic included in the Mira et al. study [49].

| Antibiotic | Class | Abbreviation |
| --- | --- | --- |
| Ampicillin | Penicillin | AMP |
| Amoxicillin | Penicillin | AM |
| Cefaclor | Cephalosporin Gen. 2 | CEC |
| Cefotaxime | Cephalosporin Gen. 3 | CTX |
| Ceftizoxime | Cephalosporin Gen. 3 | ZOX |
| Cefuroxime | Cephalosporin Gen. 2 | CXM |
| Ceftriaxone | Cephalosporin Gen. 3 | CRO |
| Amoxicillin + Clav. | Penicillin + Inhib. | AMC |
| Ceftazidime | Cephalosporin Gen. 3 | CAZ |
| Cefotetan | Cephalosporin Gen. 2 | CTT |
| Ampicillin + Sulbactam | Penicillin + Inhib. | SAM |
| Cefprozil | Cephalosporin Gen. 2 | CPR |
| Cefpodoxime | Cephalosporin Gen. 3 | CPD |
| Piperacillin + Tazobactam | Penicillin + Inhib. | TZP |
| Cefepime | Cephalosporin Gen. 4 | FEP |

**Figure 2.**
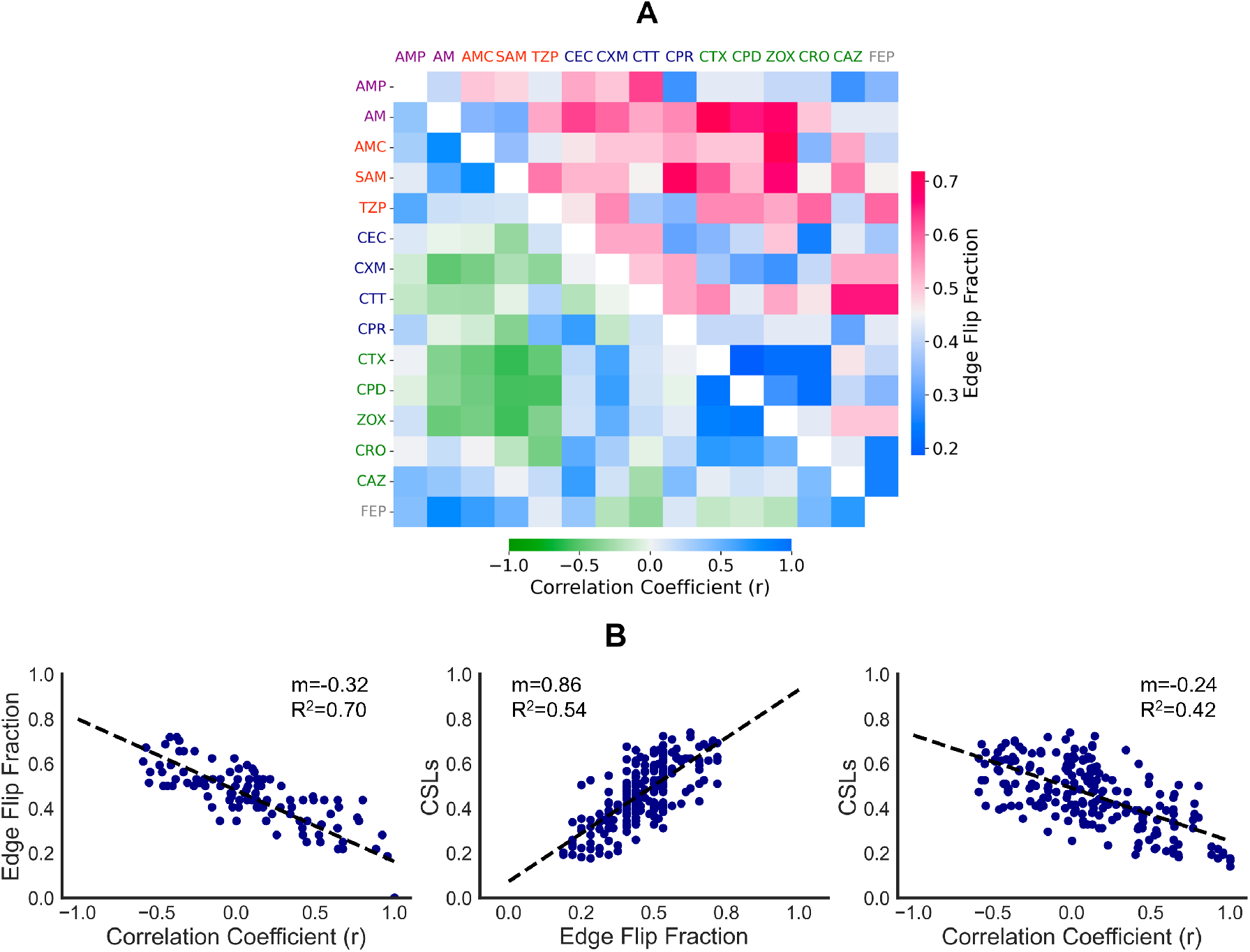
**A**. Heatmap showing the edge flip fraction (upper triangular) and the Pearson correlation coefficient (lower triangular) between pairs of empirical landscapes measure by Mira et al, with the drugs color-coded by their classes as detailed in **Table 1. B**. Scatterplots with linear fits illustrating the relationships between edge flip fraction (ℰ), Pearson correlation coefficient (*r*), and collateral sensitivity likelihood (CSLs) for the same set of empirical landscapes presented in heatmap A [49]. The slope and *R*^2^ values are indicated in the figure. Statistical significance was assessed via Pearson correlation, with p-values *<* 10^*−*60^, 10^*−*36^, and 10^*−*27^.

### Edge flip fraction correlates well with collateral sensitivity likelihoods

To go beyond descriptions of landscape topography, we explored how information derived from edge flip fraction could be utilized for rational treatment design. In particular, we examined the suitability of pairs of fitness landscapes for evolutionary control by leveraging collateral sensitivity likelihoods (CSLs) [48]. CSLs quantify the likelihood that evolution under a first drug induces susceptibility to a second. As a result, drug pairings with high CSLs are likely advantageous for therapeutic sequencing as they provoke sequential sensitivity. A direct comparison of edge flip fraction and CSLs revealed a strong positive correlation in the empirical fitness landscapes of Mira et al. (**Figure 2B**).

To further investigate the edge flip fraction in both empirical and synthetic landscapes, we analyzed pairs of fitness landscapes constructed using the rough Mount Fuji (RMF) model, Kauffman NK model and trade-off-induced fitness landscapes (TIL) model [50–53]. Despite the statistical and topographical differences between landscapes that result from a choice of the underlying model, we consistently observed an anti-correlation between edge flip fractions and Pearson correlation coefficient, as well as a positive correlation between edge flip fraction and CSLs (**Figures 3 and S2**).

**Figure 3.**
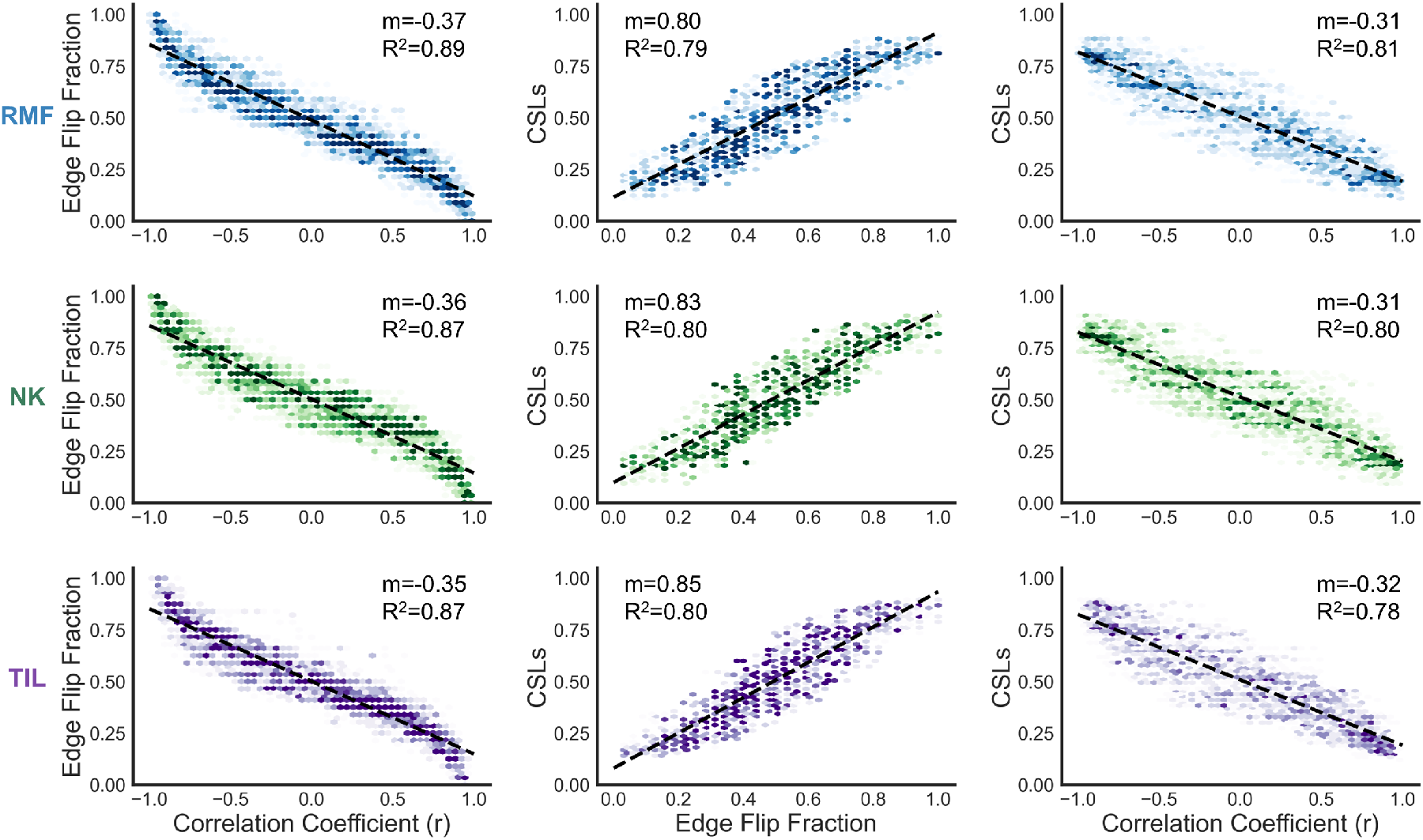
Edge flip fraction (ℰ), collateral sensitivity likelihood (CSLs), and Pearson correlation coefficient (*r*), all partially explain the variance within each other in synthetic fitness landscapes – including the rough Mount Fuji (RMF), Kauffman NK, and trade-off-induced fitness landscape (TIL) [49–53]. Hexbin plots show the density of data points, with darker regions indicating higher density, and dashed lines represent linear regression fits for the displayed relationships. As expected, ℰ and CSLs and are positively correlated, while both ℰ and *r*, CSLs and *R* are negatively correlated; all correlations are highly significant (p < 0.001).

### Edge flips enable evolutionary control in evolution on synthetic landscapes

Judicious drug sequencing has been shown promise to limit the emergence of treatment resistance in evolving populations by steering populations away from highly resistant genotypes [29, 48]. However, aside from cross resistance and collateral sensitivity, very little is known about the factors that contribute to the “controllability” of an evolving population. We hypothesized that pairs of landscapes with a high edge flip fraction are likely to be more useful for controlling evolution than landscape pairs with similar correlation coefficient, but significantly fewer edge flips. Previous evolutionary control work on fitness landscapes has shown that a Markov Decision Process (MDP) can be used to identify optimal drug policy to minimize population fitness; for any current genotype state, the MDP recommends the best drug to apply. [8]. Following this example, we compute the optimal two-drug policy to minimize population fitness for each pair of synthetically generated fitness landscapes. We then define a performance metric, *δ*, that quantifies the difference between how well the optimal policy minimized resistance, in comparison to the best single drug policy (see Methods). Intuitively, *δ* can be understood as the increase in controllability of the population fitness that is a result of optimally sequencing a pair of fitness landscapes. We observed that *δ* (controllability) grows in tandem with edge flip fraction, even for pairs of fitness landscapes with similar Pearson correlation coefficients (**Figure 4, and Figure S3**). This result can be seen most clearly (**Figure 4, right sub-panels**) when comparing two pairs of fitness landscapes each with similar correlation (*r ≈ −* 0.15) but different edge flip fractions (ℰ = 0.69 vs ℰ = 0.41). These results highlight just how critical topographical changes are to fine control of evolving populations.

**Figure 4.**
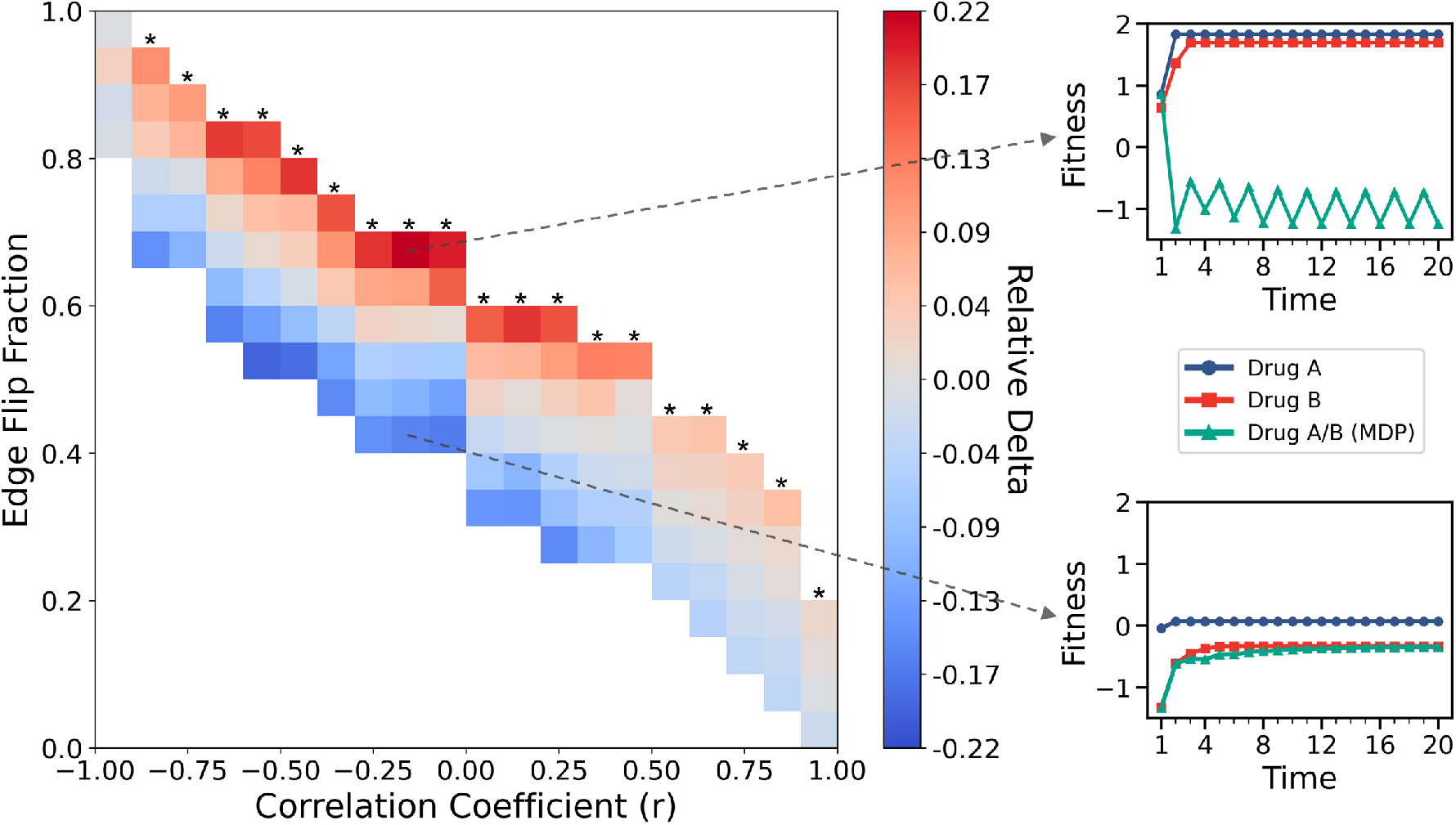
Heatmap illustrating the relationship between edge flip fraction (vertical axis) and Pearson correlation coefficient (horizontal axis) for 500,000 pairwise synthetic rough Mount Fuji (RMF) fitness landscapes. The color scale indicates the relative *δ* — that is, the improvement of the optimal two-drug MDP strategy over a single-drug strategy — normalized by the mean *δ* for each column. Asterisks (*) mark a significant trend (p < 0.001) showing that relative *δ* increases as edge flip fraction increases. The two line plots on the right compare average fitness trajectories under single-drug treatment and the optimal two-drug MDP policy in two scenarios: one with a high edge flip fraction, revealing a larger performance gap favoring MDP, and another with a low edge flip fraction, showing only an imperceptible gap.

Next, we seek to understand what role edge flips, and related metrics such as net evolutionary flow, play in determining an optimal policy. Evolutionary net flow is defined as the difference in number of evolutionary edges that point toward a genotype vs evolutionary edges that point away from a genotype, normalized by the total number of neighboring genotypes. As a result, a value of -1 indicates all edges connected to the genotype lead to fitter neighbors (a local fitness minimum), while a value of 1 indicates all edges point to the genotype from less fit neighbors (a local fitness maximum). To do so we apply our MDP framework to the 15 empirical fitness landscapes measured in Mira et al. [8, 49]. The MDP selects the drug associated with the lowest fitness among available options for 75% of genotypes (12 out of the 16), as shown in **Figure 5A**. In the remaining 25% of cases where the MDP does not select the lowest-fitness drug, 75% of these genotypes (3 out of 4) are local fitness minima (**Figure 5B**). By generating 1000 sets of synthetically generated fitness landscapes, each containing 15 fitness landscapes of *N* = 4, we further explored this behavior. Across these stimulated landscapes, an average of 71% of MDP-selected drugs were the lowest-fitness option, and 60% of non-lowest-fitness drug choices occurred at local minima. These findings suggest that when the optimal policy deviates from drugs that are the global fitness minimum, it often does so by selecting drugs that are still local minima, likely resulting in an abundance of edge flips and therefore evolutionary control.

**Figure 5.**
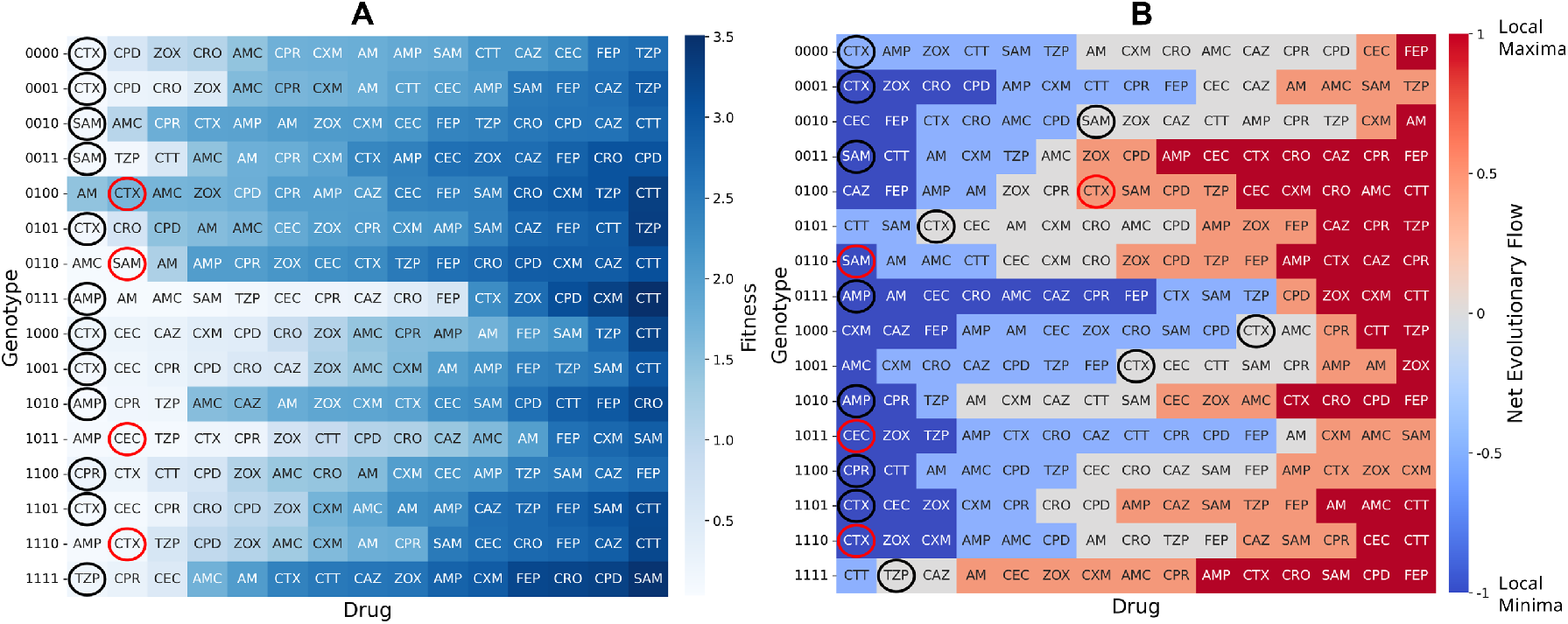
**A**. Heatmap of fitness values for 16 genotype under various drugs. The color gradient from light blue to dark blue reflects fitness levels, with darker shades representing higher fitness. Drugs selected by optimal MDP are circled: black circles denote chosen drugs with the lowest fitness, and red circles denote chosen drugs without the lowest fitness. **B**. Heatmap of the net evolutionary flow for 16 genotypes across different drugs. A value of -1 (blue) represents a local fitness minima, where all edges connected to the genotype lead to fitter neighbors, while a value of 1 (red) corresponds to a local fitness maxima, where all edges point toward the genotype from less fit neighbors. As in heatmap A, drugs selected by optimal MDP are marked with black circles for the lowest-fitness choice and red circles otherwise.

To elucidate the relationship between the edge flip fraction and the optimality of drug sequences, we simulated 1,000 drug sequences of length-20 by repeatedly applying the MDP optimal drug policy. For each drug sequence, we computed both global edge flips - occurring across the entire landscapes - and local edge flips specific to the current genotype when switching between drugs. As a contrasting baseline, we also stimulated a “worst-case” MDP policy designed to maximize, rather than minimize, population fitness over time (**Figure S4**). We then compared global and local edge flips across four sequence types: optimal drug sequences, shuffled optimal sequences (using the same drugs as the optimal sequence but in randomized order), random drug sequences (drugs chosen randomly from the full set), and the worst-case sequences (**Figure 6**).

**Figure 6.**
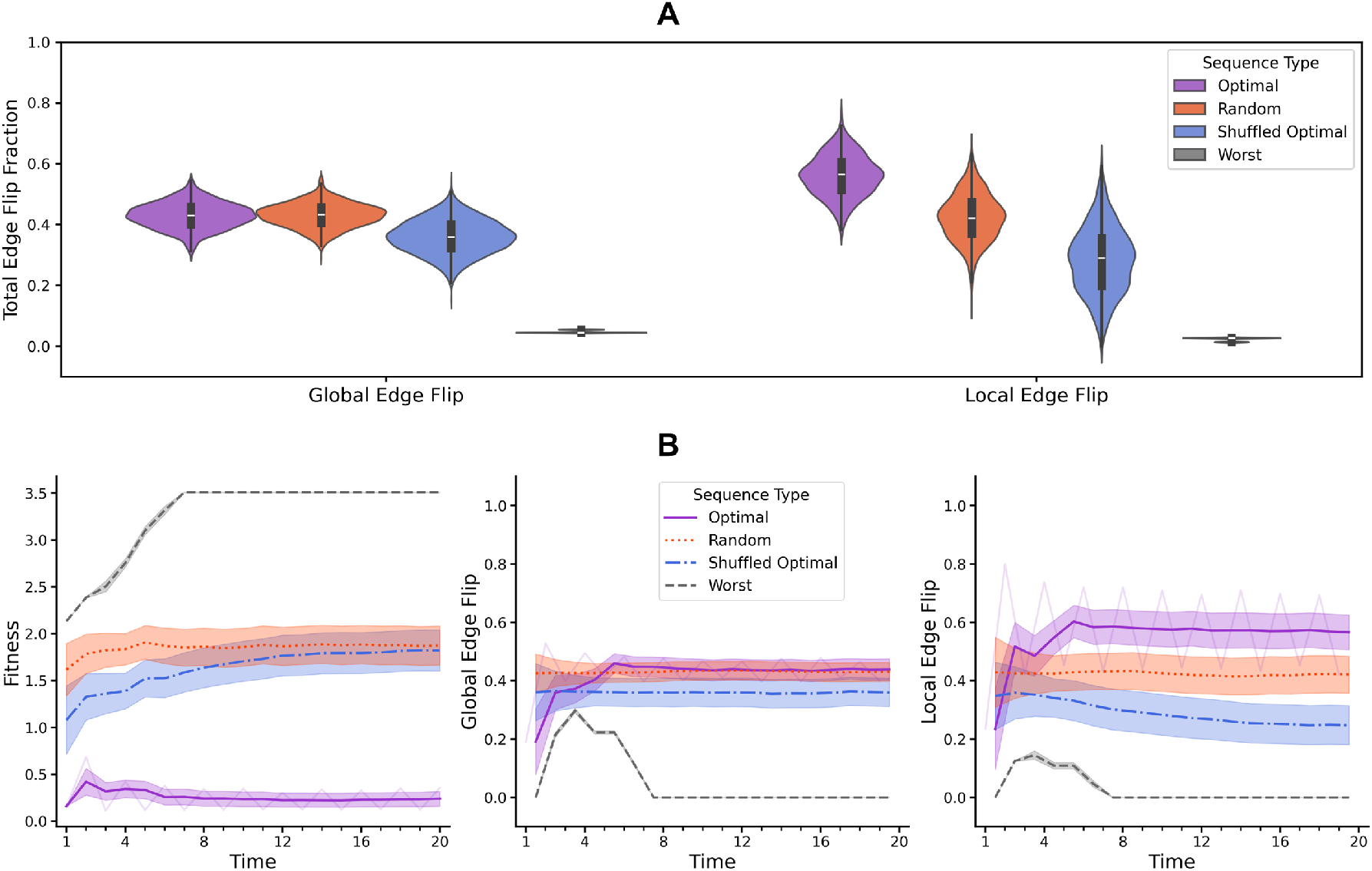
Comparison of edge flip dynamics and fitness across optimal, shuffled optimal, random and worst drug sequences. **A**. Violin plots showing the distribution of total edge flip fractions for global edge flip (across the entire landscape) and local edge flip (specific to the current genotype) across four drug sequence types: Optimal, Random, Shuffled Optimal, Random and Worst. The worst sequences show significantly lower global and local edge flip fraction compared to other sequences. While global edge flips do not differ significantly among optimal, shuffled optimal and random sequences, optimal drug sequences exhibit notably higher local edge flips compared to other sequence types. **B**. Time-series line plots tracking changes in fitness, global edge flips, and local edge flips over time. Optimal sequences (purple) maintain low fitness values with high local edge flip in cyclical dynamics. Shuffled optimal (blue) and random (orange) sequences show moderate fitness levels with moderate local edge flips. The worst sequence (gray) consistently exhibits the highest fitness and minimal edge flipping. To aid visualization of overall trends, all curves are shown as sliding-window averages (window size of 4 steps). For the optimal sequence, which displays large-amplitude periodic behavior, the original unsmoothed curve is overlaid transparently in the background. Shaded areas indicate the standard error of the mean across replicates.

The results revealed that the worst drug sequences exhibited the lowest fraction of global and local edge flip, concomitant with the highest fitness levels. This outcome arises because the worst MDP policy tends to switch drugs less often, allowing the genotype evolve toward the highest fitness peak more rapidly. While global edge flip fraction did not show significant differences among the optimal, shuffled optimal and random drug sequence - since the MDP optimizes the drug choice for each genotype rather than altering the entire fitness landscape - there were significantly more local edge flips observed in the optimal drug sequences. This phenomenon occurs because the MDP tends to select drugs that drive the genotype into a local minimum to maintain lower fitness levels. In subsequent evolutionary steps, the genotype is likely to evolve stochastically to a fitter neighbor. The optimal drugs applied thereafter tend to reverse evolutionary direction, resulting a local edge flip.

By inducing oscillations between genotypes near local minima, the optimal MDP policy effectively traps the population in less favorable states. This strategy leverages the topographical features of the fitness landscapes, exploiting the arrangement of local minima and the connectivity between genotypes to restrict adaptive pathways. The increased fraction of edge flips in the optimal drug sequences signifies a deliberate manipulation of evolutionary pathways to hinder the development of higher fitness genotypes. These findings underscore the potential of adaptive drug sequencing strategies in controlling the evolution of drug resistance.

## Discussion

Drug-resistant populations rarely evolve a single, static environment; instead, changing drugs or conditions continually reshapes their fitness landscape. To evaluate how fitness landscapes change when switching from one environment to another, we introduced the edge flip fraction. This metric measures topographical alterations by identifying what fraction of pairwise genotype relationships invert between two distinct landscapes. Specifically, an “edge flip” happens when a previously beneficial mutation becomes deleterious under the new condition, or vice versa. In essence, edge flip fraction identifies shifts in preferred evolutionary direction in a genotype network. It correlates well with established metrics like Pearson correlation coefficients and CSLs: high Pearson *r* between two landscapes implies few edge flips (and typically a low CSL, indicating cross-resistance), whereas divergent landscapes with many sign inversions yield lower correlations and higher probabilities of collateral sensitivity. Even though edge flip fraction correlates with existing measures, it provides distinct advantages in evaluating optimal drug policies. Specifically, among pairs of landscapes with comparable Pearson *r*, those featuring a higher edge flip fraction consistently yield better outcomes under an MDP-derived optimal switching protocol compared to single-drug regimens. This underscores how frequent directional changes in evolutionary pathways can significantly enhance the effectiveness of carefully coordinated therapy.

In the 15 empirical landscapes from Mira et al., MDP-derived optimal policies that sustain lower fitness levels consistently trigger more local edge flips than less-optimal approaches, effectively enforcing a directional change at each step. By switching the environment to one in which the pathogen’s current genetic state is suboptimal, the treatment is constantly forcing the population to “pay a price” for having adapted to the previous drug. Every time the population climbs a hill in one landscape, the next drug change turns that hill into a valley. Our findings align with earlier notions that sequential therapy can trap evolution in low fitness genotypes [8, 34, 48].

Our analysis, however, rests on the strong-selection, weak-mutation (SSWM) limit, where the population is effectively monomorphic; beneficial mutants arise one at a time and fix before the next mutation appears. In this limit only the sign of a fitness effect matters, so reversing that sign—our “edge flip” – completely reverses evolutionary flow. Outside SSWM, however, multiple beneficial mutants can coexist and compete (clonal interference) [54], and fixation probabilities depend on the magnitude of fitness differences, not just their sign. Consequently, two landscapes could exhibit identical edge-flip fractions yet yield different evolutionary outcomes because larger fitness advantages sweep more quickly; edge flips that merely reduce (but do not reverse) a fitness advantage can meaningfully alter trajectories; and high mutation supply can open “leap-frogging” paths that bypass flipped edges altogether [55]. These complications caution against using edge-flip fraction alone when mutation is frequent, populations are large, or horizontal gene transfer accelerates diversity. Extending or complementing the metric with information on fitness magnitudes and mutation supply will be an important next step.

While our study focused on abrupt environmental shifts (discrete changes from one drug to another), many real-world scenarios involve more continuous or gradual changes in conditions, such as fluctuating drug concentrations over time within a patient. A promising direction for future work is to extend the notion of the edge flip fraction to fitness seascapes, where the “landscape” evolves continuously based on an environmental parameter (e.g., drug dose) rather than in a stepwise fashion. Recent studies propose fitness seascapes as a more realistic model of evolution in fluctuating environments [28, 56, 57]. In scenarios where conditions differ only slightly (e.g., the same drug at different doses) rather than representing entirely separate environments, changes in edge direction between vertices of a fitness landscape can convey more nuanced information. For instance, if the fitness of two genotypes switches rank order with increasing dose of a drug, and these genotypes have mutational distance (i.e. hamming distance) of one, an edge flip must have occurred. In this way, changes in evolutionary direction between distinct conditions can relate to topographical similarity, whereas edge flips with changing of some continuous environmental variable like drug dose or temperature can be the result of a fitness cost.

In summary, inverted fitness topographies offer a compelling mechanism to steer evolutionary outcomes, and our work defines a metric and method to harness this mechanism. The insights gained are significant for providing insights into evolution and can facilitate improved evolutionary control techniques.

## Methods

### A Markov Model of Evolution on Fitness Landscapes

We begin by considering Wright’s Fitness Landscape, or a mapping of genotype and fitness. As established in previous studies [17, 48], we imagine that each of *N* loci can be either wild-type or contain a point mutation. Any given genotype is then represented by a bit-string of length *N* where bit 0 represents the absence of a point mutation, and bit 1, presence. This gives us 2^*N*^ possible genotypes which can be represented as an *N* dimensional hypercube with vertices in {0, 1}^*N*^ and edges connecting mutational neighbors. The fitness of a genotype, driven by selection from the environment, can be presented by the following fitness function:

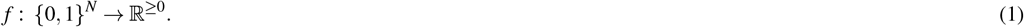

It is assumed that the entire population occupies one genotype at a time, or in other words, the population is isogenic. This assumption is valid when mutation rate *μ* is sufficiently low and selective advantage *s* is sufficiently high relative to effective population size *N*_*e*_ such that a mutation fixes on much shorter times scales than the waiting time for the next mutation – that is, when *N*_*e*_*μ* ≪1 and *N*_*e*_*s* ≫1. This regime is also known as the strong selection weak mutation (SSWM) regime, as dubbed by Gillespie [58, 59]. The details of how the population moves from genotype to genotype depending on their fitness varies with the model of evolution employed. In this paper, the population’s evolution will be described as a Markov chain governed by the random move model [48]. At each evolutionary time step, the population transitions to a fitter neighboring genotype (at Hamming distance of 1) with uniform probability, and the probability of these transitions is encoded in the transition matrix *P* = ℙ [(*i→ j*). By leveraging the properties of Markov chains, it can be shown that the population reaches a local fitness peak in finite evolutionary steps [48].

### Calculating the Edge Flip Fraction between Pairs of Fitness Landscapes

The resultant transition matrices described in the previous section define directed acyclic graphs with self-loops at the local fitness peaks. In this work, we are interested in understanding changes in the direction of evolution between pairs of fitness landscapes. For the transition matrix of two landscapes **A** and **B**, the edge flip matrix **E** is given by:

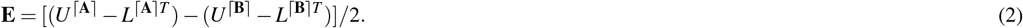

Here, *U*^⌈**A**⌉^ and *U*^⌈**B**⌉^ denote the upper triangular portions of the matrices **A** and **B**, respectively, after applying the ceiling function to each element. The ceiling function rounds each element up to the smallest integer greater than or equal to the given number, resulting in matrices consisting only of 0s and 1s. Similarly, *L*^⌈**A**⌉*T*^ and *L*^⌈**B**⌉*T*^ refer to the transposed lower triangular portions of the matrices **A** and **B**, respectively, with the ceiling function also applied to each element.

Then, the edge flip fraction, denoted by ℰ, quantifies the extent of directional changes in evolution between the two landscapes. It is calculated by summing over all elements |**e**_*ij*_| within the edge flip matrix **E** and normalizing by the total number of edges in the N dimensioinal hypercube:

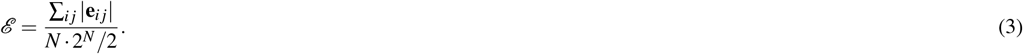

If the edge flip fraction, ℰ = 1, it indicates that all the directions of evolution differ between the two landscapes, whereas ℰ = 0 signifies no changes in the evolutionary direction. This formulation provides an analytical framework for quantifying the evolutionary direction changes between pairs of landscapes.

### Collateral Sensitivity Likelihoods (CSLs)

Collateral Sensitivity Likelihoods (CSLs), is an established metric for quantifying the probability that a population, after evolving to a local fitness peak under one drug, will exhibit increased sensitivity when subsequently exposed to a second drug [36]. Building on the Markov model described above and following established methods [36], we conducted simulations across pairs of fitness landscapes to measure CSLs.

### Construction of Paired Synthetic Fitness Landscapes

We generated synthetic fitness landscapes using a many-peaked “rough Mt. Fuji” (RMF) model of size *N* = 4 (yielding 16 total genotypes) [51, 60]. As in previous work [50], for each base landscape (denoted as A), we constructed a set of correlated landscapes (B) spanning a wide range of Pearson correlation coefficients from -1 to 1.

For the initial exploration of the edge flip fraction and its correlates (**Figure 3**), we used 50 base landscapes, each paired with 50 correlated variants (2,500 total pairs). In subsequent analyses examining both the edge flip fraction and the optimality of the drug sequence (**Figure 4**), we increased the sampling to 1,000 base landscapes and 500 correlated variants each, resulting in 500,000 total pairs and a smoother distribution of outcomes. We repeated the same procedure using the Kauffman NK model [52], producing additional sets of landscapes and correlated variants in each case.

Finally, using an empirically grounded Tradeoff-Induced Landscape (TIL) model [15], which provides genotype-specific dose-response curves (fitness seascapes), we extracted a fitness landscape at the concentration corresponding to *IC*_90_ (landscape A). We then generated an ensemble of correlated landscapes (B) with the same correlation methodology described above.

### Markov Decision Process (MDP)

We employed a Markov Decision Process (MDP) framework [8, 29, 34] to systematically identify the “optimal” drug policies that minimize population fitness. In this framework, each genotype corresponds to a discrete state. At each timestep, the system transitions stochastically to a new state according to predefined probabilities, and one of several possible actions (e.g., choosing a specific drug) is selected. The chosen action determines both an immediate reward (e.g., a change in fitness) and the probabilities governing the subsequent state transition. By iterating over all possible states and actions, the MDP computes a policy that optimizes a given objective function.

First, we applied the MDP to a pair of synthetic fitness landscapes to find an “optimal” policy that minimizes population fitness. We then compared the long-term average fitness achieved by the MDP policy, 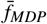, against the long-term average fitness under single-drug treatments, 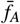 and 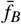. To quantify this comparison, we defined

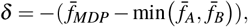

where min 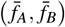 represents the lower (i.e. better) average fitness of the two single-drug outcomes. Thus, *δ* indicates how much lower the MDP-optimized fitness is relative to the best single-drug scenario, providing a direct measure of how effectively the MDP improves fitness outcomes.

Lastly, we applied the MDP to a set of 15 empirical fitness landscapes from Mira et al., where we identified not only an “optimal” policy that minimizes population fitness over time, but also a “worst” policy that maximizes population fitness.

## Supplementary Material

**Figure S1.**
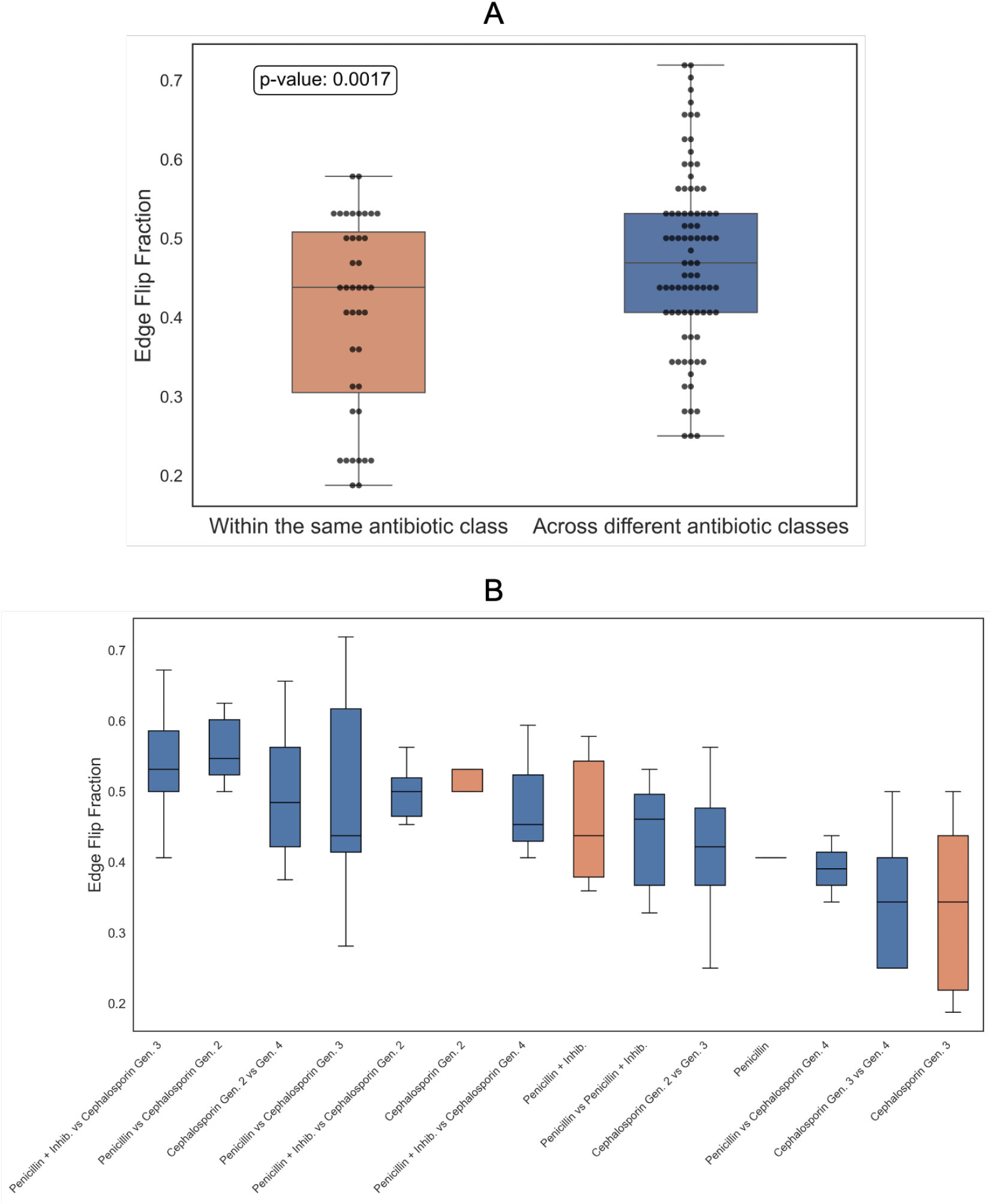
**A**. Box plot illustrating that the edge flip fraction is significantly lower in pairs of fitness landscapes from the same antibiotic class compared to those from different antibiotic classes. Antibiotic class details are provided in **Table 1. B**. Box plot displaying edge flip fractions for pairs from the same and different antibiotic classes, sorted by mean values in descending order from left to right.

**Figure S2.**
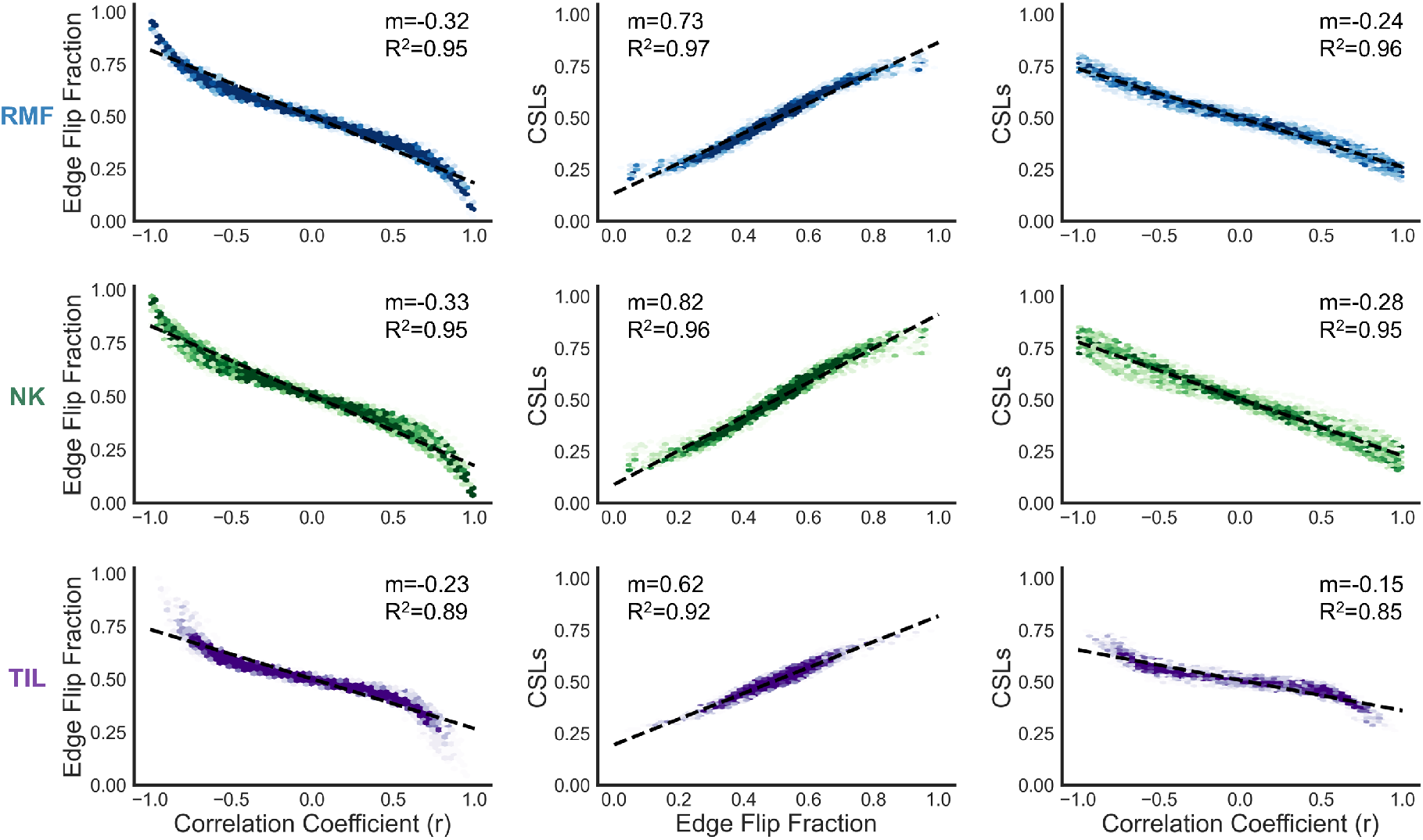
Comparison of synthetic *N* = 8 fitness landscapes generated using the rough Mount Fuji (RMF), Kauffman NK model, and trade-off-induced landscapes (TIL) [49–53]. The observed relationships - ℰ and CSLs being positively correlated, while ℰ and *r*, as well as CSLs and *R*, being negatively correlated - remain consistent in pairs of higher-dimensional fitness landscapes.

**Figure S3.**
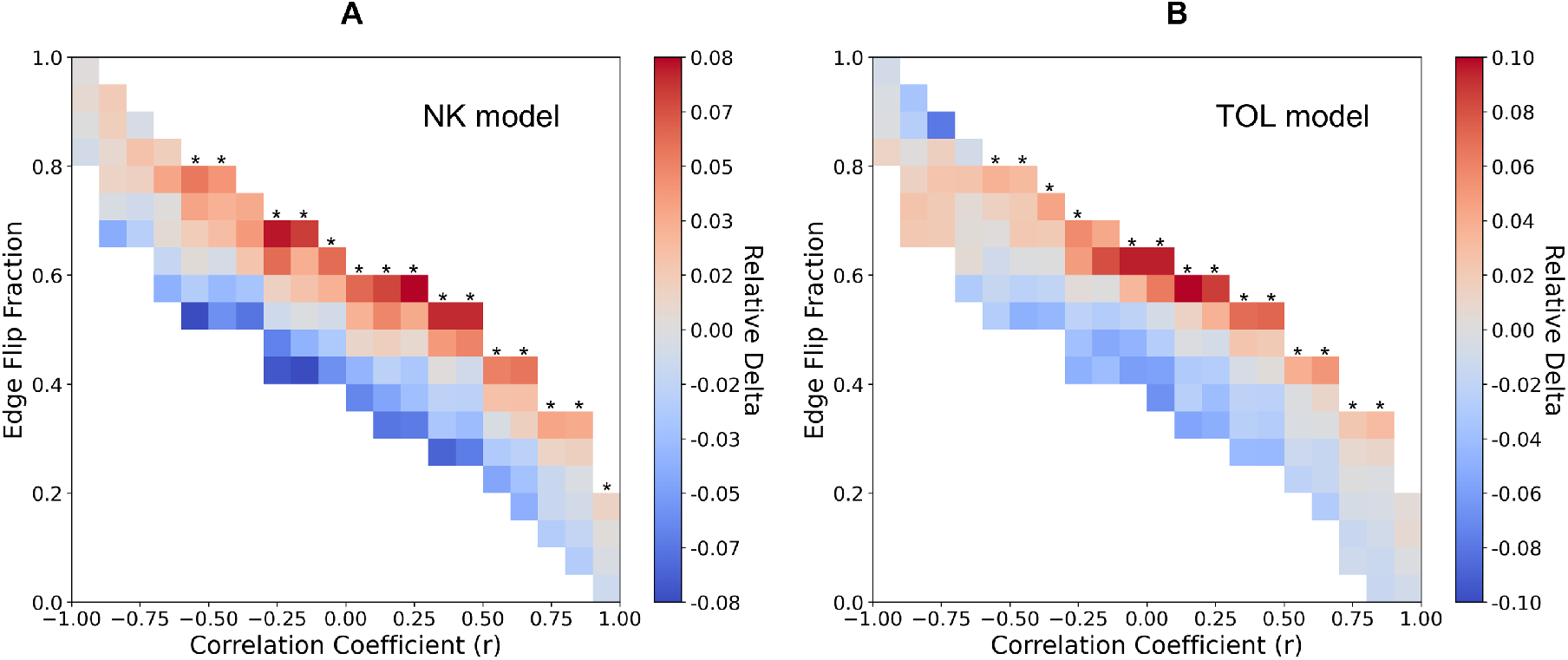
Heatmaps showing the relationship between edge flip fraction (vertical axis) and Pearson correlation coefficient (horizontal axis) for pairs of synthetic fitness landscapes generated by **A**. the Kauffman NK model and **B**. trade-off-induced landscapes (TIL). The color scale indicates the relative *δ* — that is, the improvement of the MDP-based strategy over a single-drug strategy — normalized by the mean *ε* for each column. Asterisks (*) mark a significant trend (p < 0.001) demonstrating that relative *δ* increases with higher edge flip fraction.

**Figure S4.**
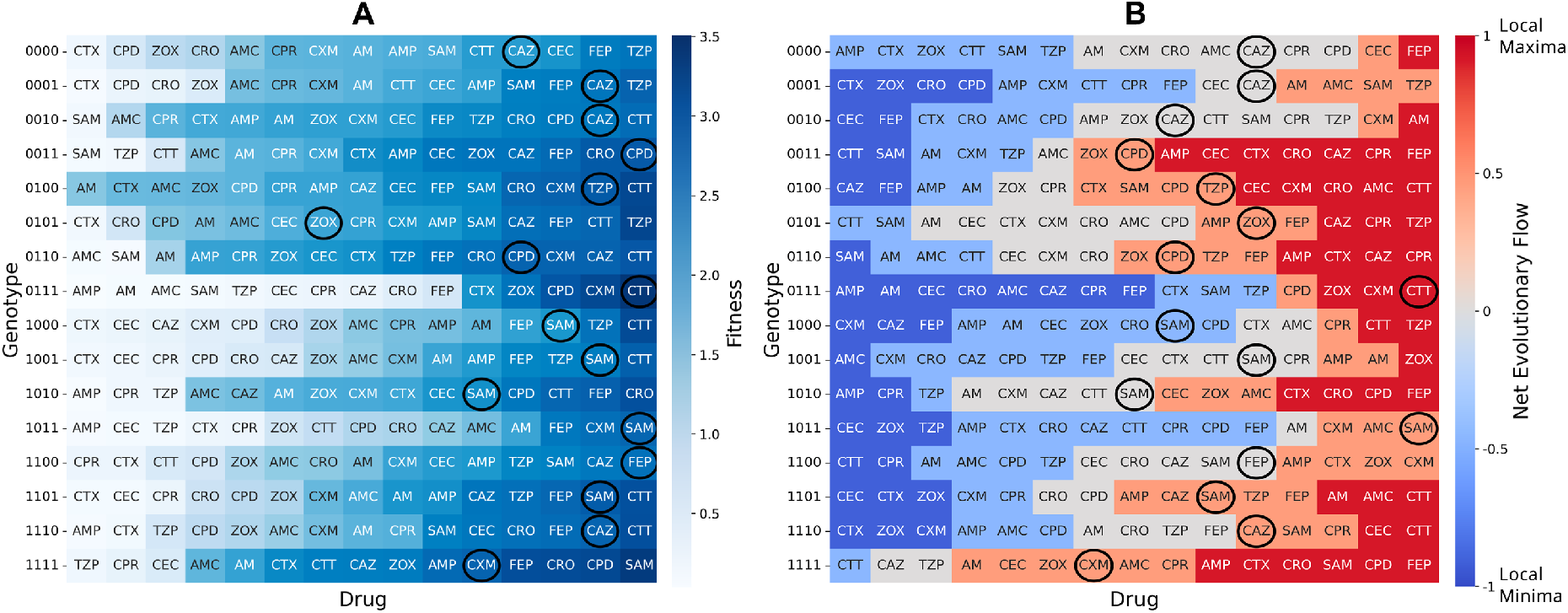
**A**. Heatmap of fitness values for 16 genotype under various drugs. The color gradient from light blue to dark blue reflects fitness levels, with darker shades representing higher fitness. Drugs chosen by the “worst-case” MDP policy for each genotype are highlighted with black circles. **B**. Heatmap of the net evolutionary flow for 16 genotypes across different drugs. A value of -1 (blue) represents a local fitness minima, where all edges connected to the genotype lead to fitter neighbors, while a value of 1 (red) corresponds to a local fitness maxima, where all edges point toward the genotype from less fit neighbors. Drugs selected by “worst-case” MDP policy are marked with circles.

